# Aquaporin-1 sustains lymphangiogenic responses in inflammatory microenvironments

**DOI:** 10.1101/2025.03.05.641652

**Authors:** Irena Roci, Jaeryung Kim, Kelly de Korodi, Tania Wyss, Jeremiah Bernier-Latmani, Silvia Arroz-Madeira, Esther Bovay, Hans Schoofs, Taija Mäkinen, Agnès Noël, Tatiana V. Petrova

## Abstract

Intestinal lymphatic vessels are essential for dietary lipid absorption and immune cell trafficking. Specialized villus lymphatic capillaries, lacteals, undergo constant VEGF-C-dependent renewal to maintain their function in a hyperosmolar, inflammatory microenvironment exposed to dietary by-products. The mechanisms of lacteal adaptation remain incompletely understood. We integrated new and published single-cell RNA-sequencing data to profile murine small-intestinal lymphatic endothelial cells (LECs) and identified three distinct subsets. Lacteal LECs display a transcriptional signature resembling *Ptx3*⁺ immune-interacting LECs characterized by high expression of *Aqp1*, encoding the aquaporin-1 water channel. LEC-specific deletion of *Aqp1* reduced lacteal length, impaired lipid uptake, and limited weight gain on a high-fat diet, underscoring the importance of water homeostasis in lacteal maintenance. AQP1 also promoted VEGF-C-dependent LEC migration under osmotic stress and, uniquely, was upregulated during inflammatory remodelling in secondary lymphedema and lymphatic malformations, but not during embryonic lymphangiogenesis. These findings link lacteal regeneration to inflammatory lymphatic remodelling and highlight tissue osmolarity as a key biophysical factor in postnatal lymphangiogenesis.

## Introduction

The lymphatic vascular system plays a crucial role in maintaining fluid homeostasis, lipid absorption, and immune surveillance (1–3). Interstitial fluid and immune cells are first taken up by blind-ended lymphatic capillaries, from which lymph is transported through valved collecting vessels to lymph nodes and eventually returned to the bloodstream. Recent single-cell RNA sequencing (scRNA-seq) studies have revealed distinct molecular profiles of capillary, pre-collecting, and valve lymphatic endothelial cells (LECs), reflecting specialized functional compartments of the lymphatic vasculature (4–7). Similarly, blood endothelial cells exhibit transcriptional heterogeneity reflecting their functional specialization into arteries, veins, and capillaries. Additionally, multiple organ-specific blood endothelial cell subsets, such as blood-brain barrier ECs, liver sinusoidal ECs, lung aerocytes, and lipid-processing endothelial cells, have been described (8,9). However, while lymph node LECs display notable specialization (6,10,11), LECs across different organs, including the small intestine, appear to be more homogeneous than blood endothelial cells (12,13).

The primary functions of small intestinal lymphatics include dietary lipid absorption and regulation of immune responses to gut microbiota and dietary antigens (14,15). Lacteals, the specialized lymphatic capillaries located in small intestinal villi, serve as the main sites of dietary lipid uptake. Unlike other adult lymphatic vessels, lacteal LECs maintain a pro-lymphangiogenic phenotype (16,17). Following absorption, lymph is transported through submucosal vessels to the mesenteric collecting vessels for systemic distribution. Beyond their role in nutrient transport, recent evidence demonstrates that intestinal LECs from crypt-associated lymphatic vessels secrete factors such as R-SPONDIN-3 and reelin, which regulate intestinal stem cells and intestinal regeneration upon injury (18–20).

Although these findings highlight intestinal lymphatics as critical for lipid uptake and tissue repair, the adaptation of lacteals to the distinct, high-osmolarity (21) and hypoxic (22) environment of small intestinal villi has yet to be fully explored. Therefore, we aimed to refine the characterization of murine intestinal LECs by integrating scRNA-seq data from new and published datasets. Our results reveal distinct transcriptional program distinguishing lacteal from submucosal and serosal intestinal LECs and show that lacteals closely resemble a recently identified subset of *Ptx3*^+^ immune-interacting LECs (5). We identify *Aqp1,* encoding a highly conserved water channel (23,24), as a member of the *Ptx3*^+^ gene signature and we demonstrate that lacteal AQP1 is necessary for dietary lipid uptake and lacteal maintenance. We also explore the regulatory mechanisms controlling AQP1 expression, including its modulation by VEGFR3 signalling and microbial cues, as well as its regulation during development and under inflammatory conditions, such as in lymphatic malformations and secondary lymphedema. Altogether, our findings connect lacteal regeneration to inflammatory remodelling of the lymphatic vasculature and emphasize the importance of tissue osmolarity in postnatal and adult lymphangiogenesis.

## Results

### Characterization of small intestinal LEC subsets

To comprehensively explore the heterogeneity of LECs in the mouse small intestine, we conducted an integrated scRNA-seq analysis by combining datasets from our current study (samples 1-5) and three published datasets (samples 6-8) (12,20,25, Figure 1A). We identified and selected 6802 LECs based on the expression of the pan-endothelial marker *Pecam1*, lymphatic markers *Prox1* and *Vegfr3* (*Flt4*), and the absence of the blood endothelial marker *Flt1* (26). Visualization of the integrated intestinal LEC datasets using Uniform Manifold Approximation and Projection (UMAP) revealed a single, continuous grouping of cells, indicating overall transcriptional similarity (Figure S1A). However, unsupervised clustering separated the cells into three subclusters, revealing underlying heterogeneity within this population (Figure 1B). Among the cluster defining markers, Cluster 0 showed elevated expression of *Aqp1*, *Fabp4*, and *Itih5*; Cluster 1 was enriched for *Clca3a1*, *Fxyd6*, and *Klhl4*; and Cluster 2 was defined by higher levels of *Isg15*, *Itih3*, and *Irf7* (Figure 1C).

**Figure 1.**
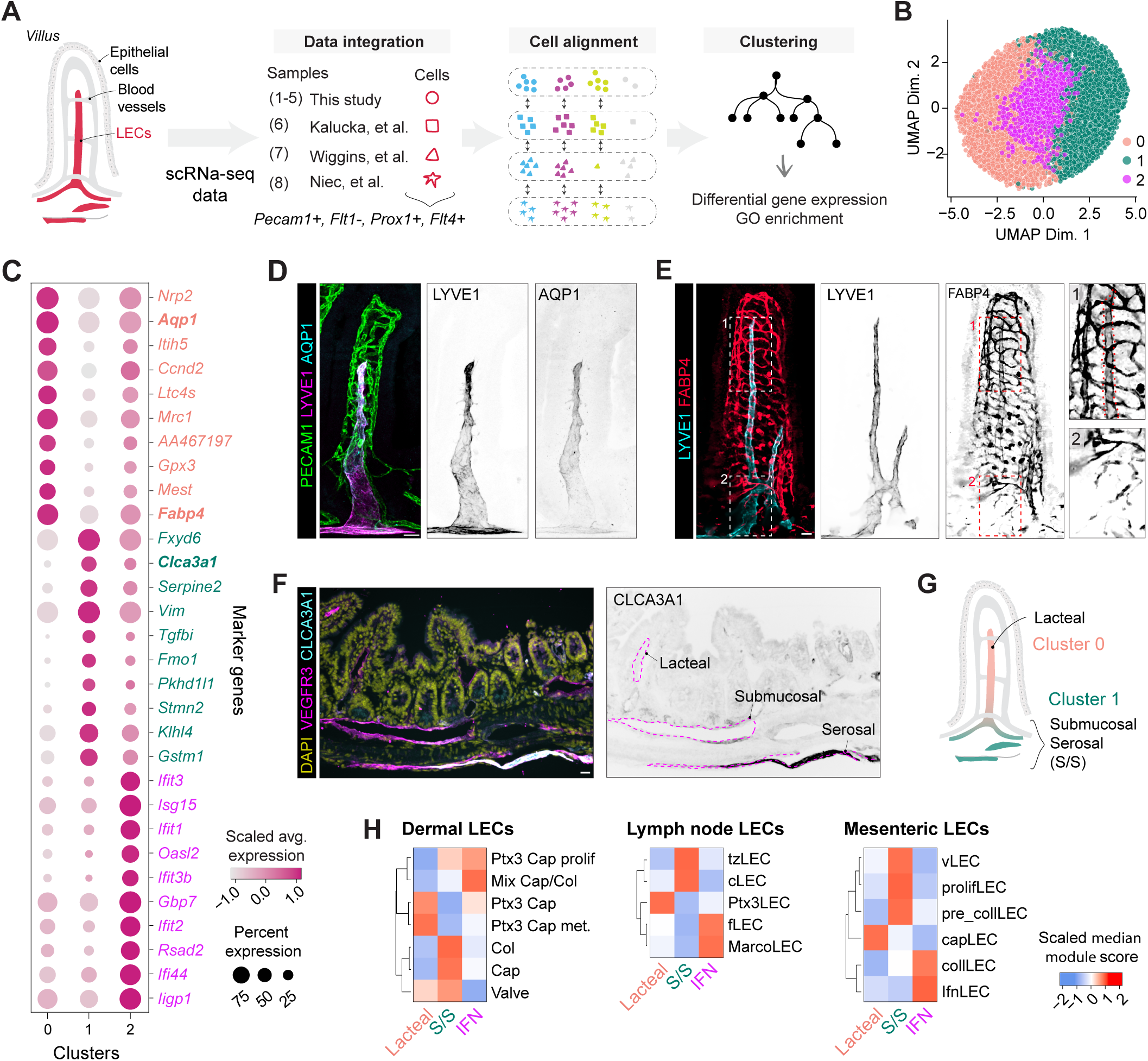
Characterization of intestinal LEC heterogeneity through integrated scRNA-seq analysis. **A.** Scheme illustrating the integration of scRNA-seq data from intestinal LECs from this study (samples 1-5) and three published studies (samples 6-8). LECs were defined as *Pecam1*^+^, *Flt1*^−^, *Prox1*^+^, and *Flt4*^+^ cells. Cells with similar transcriptional profiles were aligned and grouped to minimize batch effects, followed by unsupervised clustering, differential gene expression and GO enrichment. **B.** UMAP showing integrated cells, colored by cluster assignment at resolution 0.3. **C.** Dot plot displaying the top variably expressed genes per cluster. Dot size indicates the percentage of cells expressing the gene within each cluster; color reflects scaled average expression levels. **D.** AQP1 is expressed in lacteal LECs. Whole mount immunofluorescence staining of small intestine for PECAM1 (green), LYVE1 (magenta), and AQP1 (cyan) in the merged image. Separate grayscale images show LYVE1 or AQP1 staining alone. **E.** FABP4 is expressed in lacteal LECs. Whole mount immunofluorescence staining for LYVE1 (cyan), and FABP4 (red). Separate grayscale images show LYVE1 or FABP4 staining alone; the red dotted line in the zoomed inset outlines the lacteal. **F.** CLCA3A1 is mostly expressed in serosal LECs. Immunohistochemical staining of paraffin-embedded small intestine sections for CLCA3A1 (green), VEGFR3 (blue) and DAPI (pink). Separate grayscale image shows CLCA3A1 staining alone. **G.** The anatomical localization of the LEC clusters in small intestine, showing lacteal LECs (cluster 0), submucosal, muscular and serosal (S/S) LECs (cluster 1). **H.** Comparison of intestinal LEC clusters with LECs from other tissues. Heatmaps show the signature comparisons of three intestinal LEC clusters with LEC clusters from skin (Petkova et al., 5), lymph node (Xiang et al., 6), and mesentery (González-Loyola et al., 7), respectively. Colors represent similarity scores (scaled median module scores). Scale bars: 25 μm.

To validate scRNA-seq analyses and spatially localize the identified clusters we performed immunofluorescence staining on small intestine tissues. AQP1 was highly expressed in lacteal LECs with low expression in submucosal LECs (Figure 1D), consistent with previous findings (27–29). In addition, fatty acid-binding protein 4 (FABP4), associated with lipid transport and metabolism (30), was prominently expressed in lacteal LECs and neighbouring LYVE1^−^ blood vessels, while staining was almost absent in submucosal LECs (Figure 1E). In contrast, calcium-activated chloride channel regulator 3A-1 (CLCA3A1, formerly mCLCA1) implicated in Ca^2^⁺-dependent chloride conductance and leukocyte adhesion and migration (31–33), was strongly expressed in serosal LECs and sporadically detected in submucosal LECs, but not in lacteal LECs (Figure 1F). Based on these spatial expression patterns and prior anatomical characterizations (34), we designated Cluster 0 as lacteal LECs and Cluster 1 as submucosal and serosal (S/S) LECs (Figure 1G). Cluster 2 was characterized by increased expression of interferon-stimulated genes, suggesting a role in immune response. Both lacteal and S/S markers were also moderately expressed in these IFN cluster cells (Figure 1C), raising the possibility that upregulation of interferon signaling genes arose from tissue-processing stress. We therefore removed IFN markers (Figure S1C) and performed a new clustering analysis (Figure S1B), which yielded two clusters, with former IFN cluster cells redistributing to cluster 0 or 1. The scRNA-seq data of intestinal LECs from Wiggins et al. (25) incorporated into our bioinformatic analysis (Figure 1A, S1A) identified four LEC clusters (LEC1, LEC2a, LEC2b, and IFN), with only the IFN cluster showing transcriptional similarity to our IFN LECs. However, the spatial localization of the LEC clusters remains to be determined. Differences from Wiggins et al. likely stem from methodological factors, including LEC-only versus LEC/BEC clustering, different input parameters to the clustering algorithm (resolution and principal components), sample sizes (839 versus 6802 cells), and total number of mice analyzed (1 versus 8). Taken together, these results reveal defining transcriptional profiles and spatial organization among intestinal LECs subpopulations.

Building on this stratification, we performed additional bioinformatics analyses to characterize and predict the functional roles of these gut LEC subsets. Overrepresentation analysis of Gene Ontology (GO) pathways revealed distinct pathway enrichment among the intestinal LEC subpopulations (Figure S1C). Lacteal LECs were enriched in gene sets associated with cytoskeleton organization, growth factor responses and cell migration, consistent with previous studies showing continuous lacteal regeneration (16,17). S/S LECs showed enrichment for processes involved in RNA processing and translation, suggesting a need for increased protein synthesis. Exposure to higher mechanical stress in the submucosal and muscularis layers (35) potentially drives the need for enhanced protein synthesis to preserve vessel integrity. As expected, IFN LECs were enriched for pathways associated with immune responses and viral defence mechanisms.

In addition to analyzing gene expression, we employed SCENIC, a bioinformatics pipeline leveraging motif enrichment, to evaluate transcription factor activity in the LEC subsets (36). Lacteal LECs displayed higher activity of SOX4, MXD4, NR2F2 (COUP-TFII), HEYL, IRF8, and HMG20b, although only *Sox4*, *Mxd4*, and *Nr2f2* transcripts were clearly detected in lacteal LECs (Figure S1D, S1E). NR2F2 is essential for initiating and maintaining *Prox1* expression during LEC specification (37), while SOX4 and MXD4 have not been previously linked to lymphatic development or function. S/S LECs were enriched in regulons regulated by FOXC1, FOXC2, FOXP1, FOXP2, TBX1, EBF1, and FOXN3 (Figure S1D), while expression of *Foxp1*, *Foxp2*, *Tbx1*, and *Ebf1* was also higher in this subset (Figure S1D). Interestingly, FOXC1/2, FOXP2, and TBX1 control collecting lymphatic vessel development and function (38–40), suggesting a transcriptional similarity of S/S LECs with pre-collecting and collecting lymphatics. Finally, IFN LECs displayed enrichment of regulons for key interferon response transcription factors (Figure S1D, S1E) (41). Overall, these analyses confirm the specialized identities of the gut LEC subsets.

All gut LECs express LYVE1 and lack SMC coverage suggesting they are lymphatic capillaries by the current definitions (42). However, the above analyses suggest that lacteal and S/S LECs display distinct transcriptional characteristics. To evaluate differences between transcriptomes of intestinal LECs and those of other organs, we examined cluster-specific markers of intestinal LECs with those in dermal (5), lymph node (6), and mesenteric LECs (7) (Figure 1H). S/S LECs most closely resembled both collecting and capillary dermal LECs; pre-collecting, valve, and proliferating mesenteric LECs; and ceiling lymph node LECs (Figure 1H), which originate from afferent collecting lymphatic vessels (43). These findings reinforce the idea that submucosal lymphatics are pre-collectors. In contrast, lacteal LECs matched to capillary LECs from the mesentery, but showed an even closer alignment with *Ptx3*^+^ capillary LECs from inflamed skin and non-inflammed LN (5,6), rather than non-inflammed capillaries from these organs (Figure 1H). The lacteal signature encompassed genes specific to *Ptx3*^+^ LECs including *Itih5, Mrc1, Igfbp4, Aqp1, Ccnd2, Stab1, Mest, Tspan18*. *Mrc1* and *Aqp1* were also identified as top markers of *Ptx3+* lung LECs (44). Notably, *Ptx3* was expressed in lacteal LECs, although it did not appear among the top genes because its expression was confined to a minor subset (likely tip LECs, Figure S1F), which can be overlooked in scRNA-seq analysis (45, 6). Altogether, these results show that intestinal LECs comprise two main populations: 1) lacteal LECs in villi, which are transcriptionally similar to *Ptx3^+^*immune-interacting LECs, and 2) S/S LECs that display transcriptional signatures of pre-collector lymphatics.

### AQP1 promotes lacteal maintenance and dietary lipid uptake

Among genes enriched in lacteal LECs, *Aqp1* ranked as one of the most defining markers (Figure 1C). AQP1 is a tetrameric integral membrane protein in which each monomer serves as an osmotically driven, water-selective pore (23). Although AQP1 has been extensively studied in the kidney, it is also expressed in multiple other organs (24). Notably, it is found in blood endothelial cells, where it regulates angiogenic sprouting, cell migration, and vascular permeability, particularly in response to osmotic stress (46, 47). Lacteals must sustain their pro-lymphangiogenic phenotype (16, 17) in an osmotically challenging environment (21). Given the established role of AQP1 in angiogenesis and its exclusive expression among aquaporins in lacteal LECs, we hypothesized that AQP1 is essential for lacteal function. Consistently, AQP1 was expressed in lacteals throughout all regions of the small intestine (Figure S2A, S2B), with significantly higher levels at the lacteal tip compared to the base (Figure 2A).

**Figure 2.**
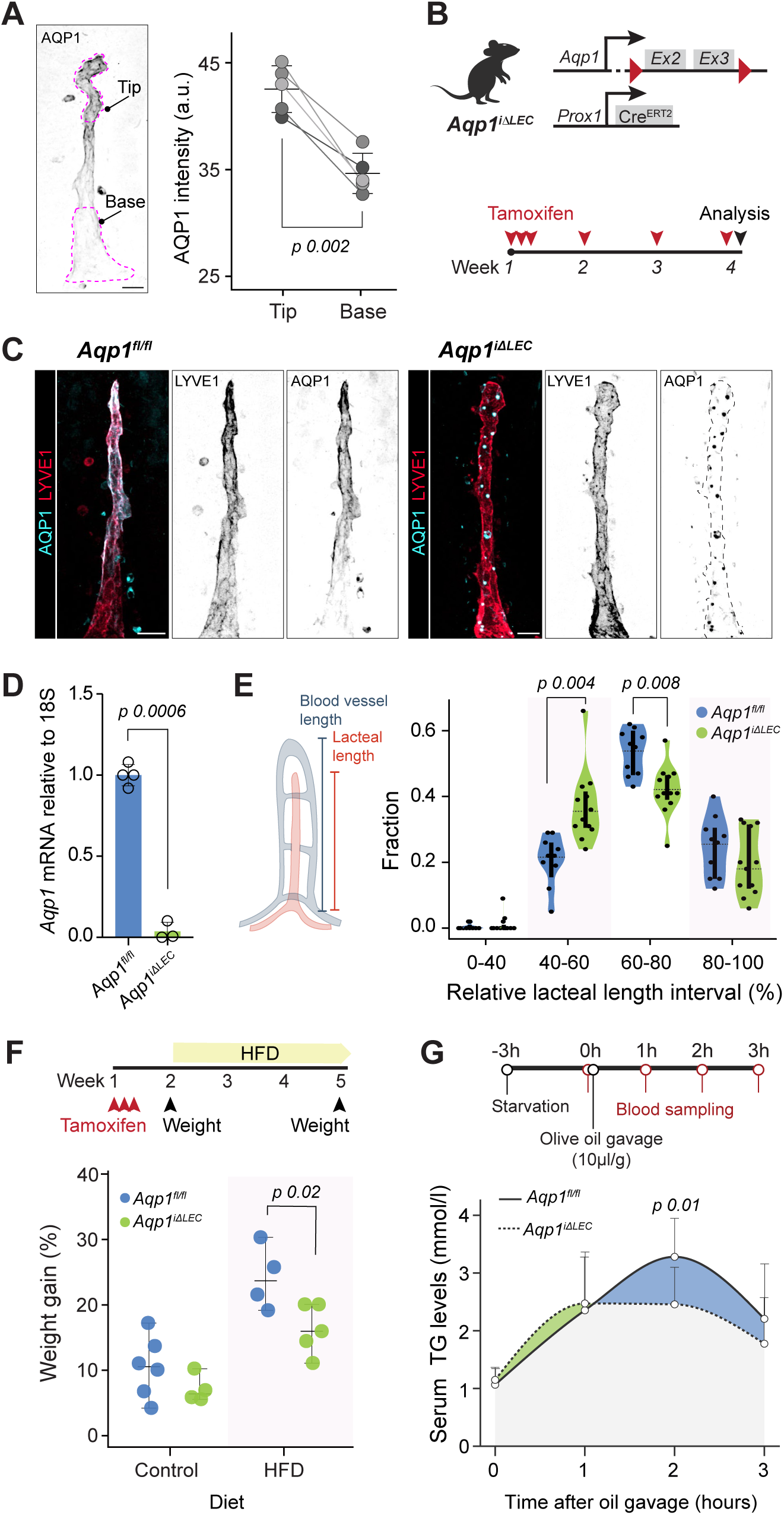
AQP1 is essential for lacteal function. **A.** AQP1 is enriched at the lacteal tip. Image of a lacteal marking the tip and base. Paired AQP1 fluorescence intensity quantification at the tip and base of the same lacteal. Differences were assessed using a paired t-test (n=5 mice, 10 lacteals per mouse). **B.** Generation of LEC-specific *Aqp1* knockout mice. Top: Schematic of the genetic model for conditional *Aqp1* deletion in LECs (*Aqp1^iΔLEC^*), with loxP sites flanking exons 2 and 3 of *Aqp1* and CreERT2 expression under the *Prox1* promoter. Bottom: Timeline of tamoxifen injections over four weeks to induce deletion. **C.** Validation of AQP1 deletion in intestinal LECs. Staining for AQP1 (cyan) and LYVE1 (red) in *Aqp1^fl/fl^* (n=4 mice) and *Aqp1^iΔLEC^* mice (n=3). Grayscale images show individual staining. Differences were assessed using a one-sample t-test. Scale bar: 25 μm. **D.** Further validation of *Aqp1* deletion. qPCR for *Aqp1* mRNA in *Aqp1^fl/fl^* and *Aqp1^iΔLEC^* mice, normalized to 18S rRNA. **E.** Lymphatic *Aqp1* deletion leads to lacteal shortening. Schematic illustrates measurement of lacteal length relative to blood vessels. Violin plots show the distribution of relative lacteal length intervals as percentages of villus length in *Aqp1^fl/fl^*and *Aqp1^iΔLEC^* mice (n=10 mice, mixed sex). Differences were assessed using a two-way repeated-measures ANOVA, followed by pairwise t-tests with Benjamini-Hochberg correction. **F.** Lymphatic *Aqp1* deletion prevents weigh gain on HFD. Timeline for tamoxifen injections and HFD administration. Dot plot shows percentage weight gain from baseline (week two) after three weeks on HFD. Differences were assessed using Welsh’s t-test for each diet (control: *Aqp1^fl/fl^*, n=6 and *Aqp1^iΔLEC^*, n=4; HFD: *Aqp1^fl/fl^*, n=4 and *Aqp1^iΔLEC^* n=5 mice). **G.** Lymphatic *Aqp1* deletion blunts dietary lipid absorption. Timeline for olive oil gavage and blood sampling at indicated intervals. Line graph shows serum triglyceride (TG) levels over time. Differences were assessed using a two-way repeated-measures ANOVA, followed by two-sample t-tests with Holm–Sidak corrections for multiple comparisons (n=11 mice). Data are shown as mean ±SD.

To directly assess the functional role of AQP1 in lacteal physiology, we generated a conditional knockout mouse model (*Aqp1^iΔLEC^*) by crossing *Aqp1^fl/fl^* mice (48), in which exons 2 and 3 of *Aqp1* are flanked by loxP sites, with *Prox1-CreERT2* mice (49), enabling tamoxifen-inducible LEC-specific deletion of *Aqp1* (Figure 2B). At the protein level, immunofluorescence staining showed a marked reduction of AQP1 in lacteal LECs of *Aqp1*^iΔLEC^ mice (Figure 2C), validating the deletion. Scattered AQP1-positive puncta 3-4 μm in size appeared along the lacteals of *Aqp1^iΔLEC^* mice, although their origin and significance remain unclear. We also confirmed efficient *Aqp1* deletion at the mRNA level by isolating LECs from the small intestine using fluorescence-activated cell sorting (FACS) and performing quantitative PCR targeting exons 2 and 3 of *Aqp1* (Figure 2D, Figure S2C).

To assess the impact of *Aqp1* deletion on lacteal morphology, we measured the relative lacteal length, calculated as the ratio of lacteal length to blood vessel length within the same villus, in *Aqp1^iΔLEC^* and control mice. The distribution of relative lacteal length intervals revealed a significant shift toward shorter lacteals in *Aqp1^iΔLEC^* mice, with an increased frequency of lacteals in the 40-60% relative lacteal length interval and a decreased frequency in the 60-80% interval (Figure 2E). These findings indicate that *Aqp1* is essential for maintaining normal lacteal length.

Given the essential role of lacteals in lipid uptake, we investigated whether *Aqp1* deletion affects lipid absorption. Mice were administered tamoxifen to induce *Aqp1* deletion and then subjected to a high-fat diet (HFD) for three weeks (Figure 2F). Under standard diet, body weight was comparable between control and *Aqp1^iΔLEC^*mice. However, under HFD, *Aqp1^iΔLEC^* mice exhibited significantly reduced weight gain compared to controls (Figure 2F), suggesting compromised lipid absorption. To directly evaluate lipid absorption, we measured serum triglyceride levels over time following oral administration of a bolus of olive oil to *Aqp1^iΔLEC^* or control mice (Figure 2G). *Aqp1^iΔLEC^* mice showed significantly lower serum triglyceride levels at two hours post-gavage compared to controls (Figure 2G), confirming impaired uptake of dietary lipids. In summary, our findings demonstrate that *Aqp1* is selectively expressed in lacteal LECs, where it contributes to lacteal maintenance and facilitates lipid absorption, highlighting the importance of water homeostasis in intestinal lymphatic physiology.

### AQP1 expression in lacteal LECs is regulated by microbiota and VEGFR3 signalling

To elucidate the mechanisms regulating *Aqp1* expression in lacteal LECs, we investigated its temporal expression during early postnatal development when lymphatic vessels actively sprout and migrate in a VEGF-C/VEGFR3-dependent manner to form lacteals (50,51). At P1 AQP1 expression in lacteals was low, sporadic and predominantly confined to the lacteal base (Figure 3A). As development progressed, AQP1 expression extended along the lacteal, with a marked increase at the tip by P5, resembling the distribution observed in adult mice (Figure 2A). Quantitative fluorescence intensity analysis confirmed this postnatal upregulation and spatial redistribution of AQP1 (Figure 3A). These findings suggest that AQP1 expression in lacteal LECs is closely linked to postnatal intestinal maturation, possibly reflecting enhanced functional requirements for lipid absorption as the neonatal gut transitions to enteral feeding (52). This transition coincides with the onset of intestinal colonization at birth, when the bacterial load increases dramatically in the first days of the postnatal period (53). Therefore, we next assessed whether microbial signals modulate AQP1 expression. We found that AQP1 expression in lacteals was significantly reduced in mice treated with broad-spectrum antibiotics compared to untreated controls (Figure 3B), while lymphatic LYVE1 staining was not changed. This data indicates that microbial cues directly or indirectly contribute to the regulation of AQP1 expression in lacteal LECs.

**Figure 3.**
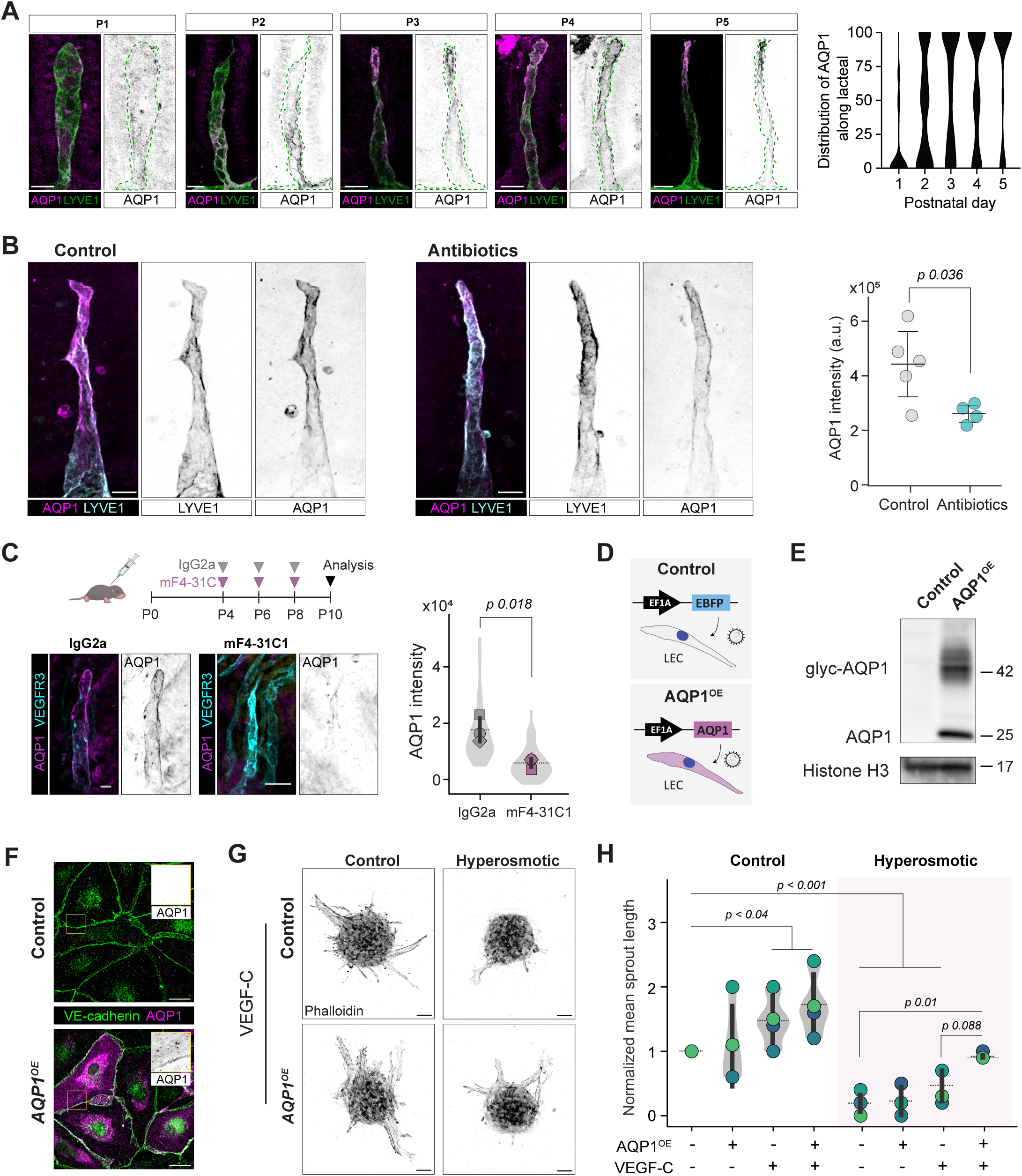
Postnatal AQP1 expression in lacteal LECs is induced by microbiota and VEGFR3 signalling. **A.** AQP1 expression emerges postnatally in lacteals. Staining for AQP1 (magenta) and LYVE1 (green) in lacteals from postnatal days P1-P5. Grayscale images show AQP1 alone. Right: Quantification of AQP1 distribution along the lacteal. One mouse per timepoint: P1 (n = 53 lacteals), P2 (n = 51), P3 (n = 55), P4 (n = 5), P5 (n = 68). Scale bar: 20 μm. **B.** Microbiota depletion reduces AQP1 in lacteals. Immunofluorescence images of lacteals from control and antibiotic-treated mice stained for AQP1 (magenta) and LYVE1 (cyan). Right: Quantification of AQP1 intensity per lacteal. Differences were assessed using Student’s t-test (control: n=5 mice; antibiotic: n=4). Scale bar: 25 μm. **C.** VEGFR3 blockade lower AQP1 expression. Timeline for administration of VEGFR3-blocking (mF4-31C1) or control IgG2a antibody. Staining for AQP1 (magenta) and VEGFR3 (cyan). Right: Quantification of AQP1 intensity per lacteal. Quantification of AQP1 intensity per lacteal. Differences were assessed using Student’s t-test (n=3 mice). **D.** Lentiviral constructs used to transduce human LECs. **E.** Validation of AQP1 overexpression in vitro. Western blot analysis of control and *AQP1^OE^*LEC lysates for the indicated proteins. **F.** AQP1 localization in transduced LECs. Staining for for VE-Cadherin (green) and AQP1 (magenta). The yellow insets show magnified regions, with AQP1 presented in grayscale. Scale bars: 25 μm. **G.** AQP1 overexpression promotes lymphangiogenic sprouting under osmotic stress. Representative grayscale images of phalloidin-stained spheroids in a sprouting assay. Scale bar: 50 μm. **H.** Quantification of sprouting. Violin plots show normalized mean sprout length under eight conditions combining cell lines (control or *AQP1^O^*^E^), VEGF-C (− or +), and medium (control or hyperosmotic). To assess genotype effects, we performed two-way ANOVAs for each medium. Under control, only growth factor was significant (p = 0.031), whereas under hyperosmotic, both genotype (p = 0.049) and growth factor (p = 0.024) were significant. P values from paired comparisons are shown in the plot. Data are shown as mean ±SD.

Lymphangiogenesis mediated by VEGF-C/VEGFR3 signalling drives the postnatal expansion of lacteals and their maintenance in adults (16,17,50). Therefore, we investigated whether VEGFR3 signalling directly regulates AQP1 expression in lacteal LECs. We administered control IgG or VEGFR3-blocking antibodies (mF4-31C1, 54) to mice at P4, P6, and P8 and analyzed AQP1 expression at P10 (Figure 3C). As expected, the treatment stunted postnatal lacteal expansion. In addition, mice treated with mF4-31C1 exhibited a significant reduction in AQP1 expression in remaining lacteals compared to IgG2a-treated controls, while VEGFR3 staining increased (Figure 3C), likely due to decreased receptor internalization (55). Collectively, these findings demonstrate that both microbial colonization and VEGFR3 signalling contribute to AQP1 expression in lacteals.

### AQP1 enhances VEGF-C-induced LEC migration under osmotic stress

Since AQP1 is enriched at the lacteal tip where osmolarity fluctuates during nutrient absorption (21), we hypothesized that AQP1 serves as an adaptation to these unique villus tip conditions. To test this hypothesis, we generated *AQP1*-overexpressing (*AQP1^OE^*) and control LECs expressing enhanced blue fluorescent protein (*EBFP^OE^*) via lentiviral transduction (Figure 3D). Western blot analysis confirmed overexpression of AQP1, displaying distinct bands corresponding to glycosylated and non-glycosylated forms (56), while control cells lacked detectable AQP1 expression (Figure 3E). Immunofluorescence microscopy further validated AQP1 expression, showing prominent membrane, junctional and cytoplasmic localization in *AQP1^OE^* LECs (Figure 3F).

To assess the impact of AQP1 overexpression on lymphangiogenic responses, we employed a spheroid sprouting assay (57). LEC spheroids were embedded in fibrin gel and cultured in the presence or absence of VEGF-C in normosmotic (∼300 mOsm) or hyperosmotic (∼500 mOsm) conditions, reflecting osmolarity observed at the villus tip (17) (Figure S3A). Under normosmotic conditions, baseline sprouting did not differ significantly between *AQP1^OE^* and control cells. However, in the presence of the key lymphangiogenic factor VEGF-C (58), both cell lines exhibited increased sprout lengths (Figure 3G). Exposure to hyperosmotic medium strongly impaired overall sprouting in both control and *AQP1^OE^* LECs. Nonetheless, *AQP1^OE^* cells retained longer sprouts than control cells when stimulated with VEGF-C (Figure 3G). LECs exposed to hypoosmotic conditions showed a tendency towards increased sprouting even without VEGF-C, although AQP1 overexpression did not affect sprouting under these conditions (Figure S3B). These data demonstrate that high osmolarity suppresses lymphangiogenic responses, even in the presence of high levels of VEGF-C, and AQP1 overexpression provides a protective advantage under hyperosmotic conditions.

### Inflammatory remodelling triggers AQP1 upregulation in adult LECs

Signalling through the VEGF-C/VEGFR3 axis activates the PI3K-AKT pathway, promoting processes essential for lymphangiogenesis and vascular remodelling, including LEC survival, proliferation, and migration (59). Activating mutations in *PIK3CA* underly development of lymphatic vascular malformations in humans (60–62). Abnormal dermal lymphatic vascular growth in *Pik3ca^H1047R^* mutant mice is maintained by accumulation of VEGF-C-producing macrophages, which selectively expand *Ptx3*-high immune-interacting LECs by paracrine signalling (63, 5). Interestingly, *Aqp1*-high lacteals share transcriptional similarities with *Ptx3*-high dermal capillary LECs (Figure 1H), and AQP1 expression partially depends on VEGFR3 signalling (Figure 3C). We therefore investigated *Aqp1* expression in dermal LECs from the *Pik3ca^H1047R^* model, by reanalyzing scRNA-seq data from the dermal LECs of *Pik3ca*^H1047R^*; Cdh5-CreERT2* mice (5) and focusing on *Aqp1* expression in LEC subpopulations. In mutant samples, *Ptx3^+^* capillary LECs were enriched and exhibited a higher proportion of cells within the highest quartile of *Aqp1* expression (75–100%) compared to capillary LECs in both control and mutant conditions (Figure 4A). Next, we explored AQP1 expression at the protein level in ears from control and *Pik3ca^H1047R^; Vegfr3-CreERT2* mice. While AQP1 expression was absent in lymphatic vessels of control ears, it was markedly increased in the hypersprouting VEGFR3^+^ LECs of mutant samples (Figure 4B). To determine whether AQP1 is also induced in VEGFR3-driven dermal lymphatic vessels during embryonic development, we examined dermal skin at embryonic day 16.5 (E16.5). AQP1 was expressed in PECAM1^+^ blood vessels but not in VEGFR3^+^ sprouting lymphatic vessels (Figure S4A). Additionally, AQP1 staining was observed in structures consistent with peripheral nerves, aligning with known neuronal expression of AQP1 (64). These results indicate that AQP1 upregulation may be specific to the PTX3^+^-capillary LEC subpopulation, arising during inflammatory lymphangiogenesis. In line with AQP1 expression in dermal *Ptx3*^+^ LECs, medullary sinus LECs in normal and inflamed lymph node, also defined by the *Ptx3*^+^ gene signature (6, Figure 1H), showed prominent AQP1 expression (Figure 4C). In contrast, AQP1 expression was either low or undetectable in subcapsular sinus LECs under both control and inflammatory conditions (Figure S4B), as well as in embryonic LN LECs (Figure S4C).

**Figure 4.**
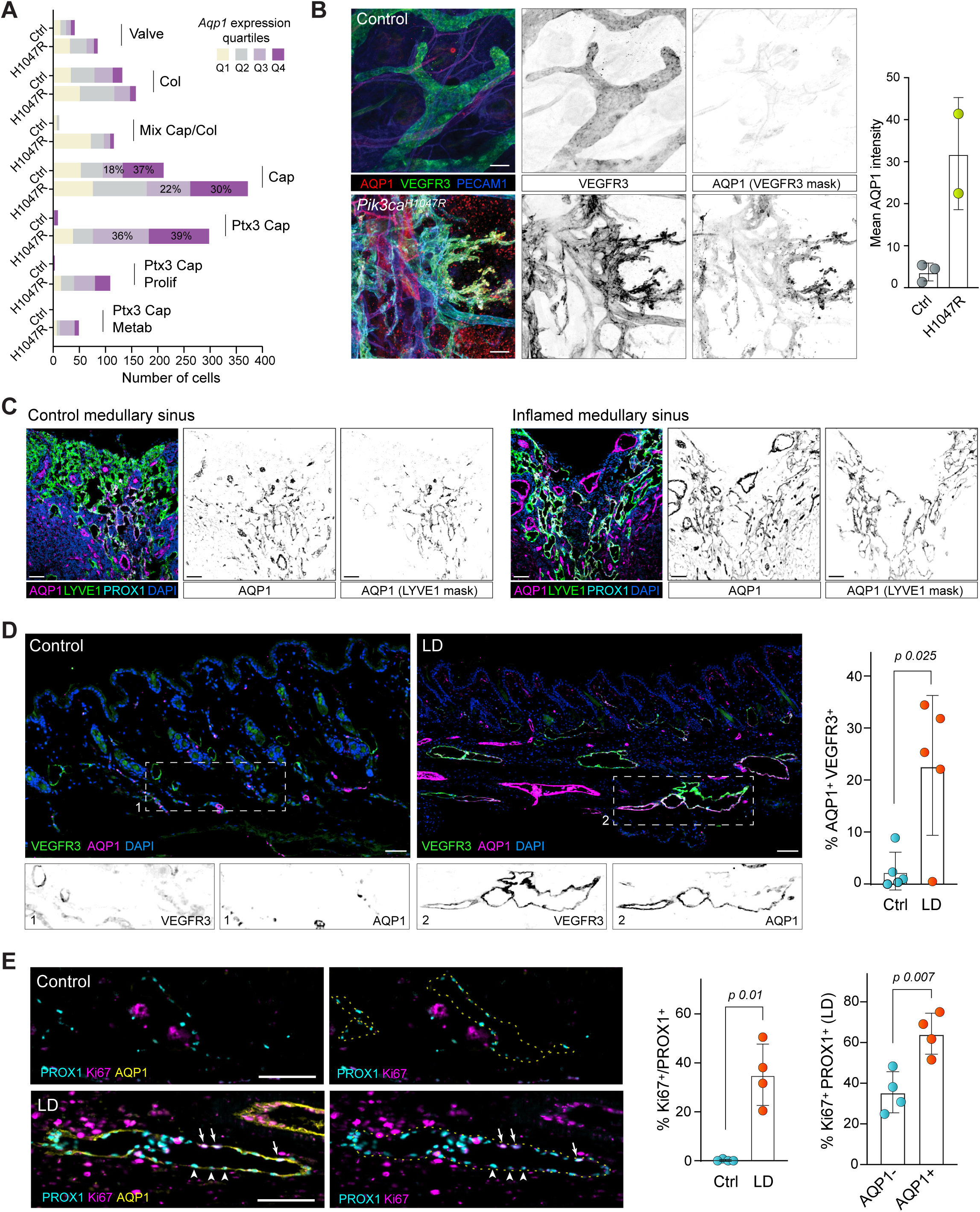
Enhanced AQP1 expression in inflammatory lymphangiogenesis. **A.** *Aqp1* levels in LEC subpopulations from control and *Pik3ca*^H1047R^ miceö RNA-seq data re-analysed from Petkova et al, (5). Colors indicate *Aqp1* expression quartiles: 0–25%, 25–50%, 50–75%, 75–100%. **B.** AQP1 is induced in hypersprouting dermal lymphatics of *Pik3ca^H1047R^* mice. Whole mount ear staining (control and *Pik3ca^H1047R^*) for AQP1 (red), VEGFR3 (green), and PECAM1 (blue). Grayscale images show individual channels. Scale bar: 50 µm. The dot plot quantifies AQP1 intensity in VEGFR3^+^ areas (control: n=3; mutant: 2 mice). **C.** AQP1 expression in medullary LN LECs. Staining of medullary sinus LECs (control and inflamed LNs) for AQP1 (magenta), LYVE1 (green), PROX1 (cyan) and DAPI (blue). Scale bar: 50 µm. **D.** AQP1 is upregulated in secondary lymphedema (LD) model. Staining of skin from control and LD mice for VEGFR3 (green), AQP1 (magenta), and DAPI (blue). Dashed squares (1: control; 2: LD) indicate insets with separate grayscale images for VEGFR3 and AQP1. Scale bar: 50 μm. The dot plot quantifies the percentage of AQP1^+^ area within VEGFR3^+^ vessels (n = 5 mice). **E.** AQP1^+^ LECs show increased proliferation. Skin sections (control and LD) stained for AQP1 (yellow), Ki67 (magenta) and PROX1 (cyan) or Ki67 (magenta), and PROX1 (cyan). Arrows indicate PROX1^+^ Ki67^+^ cells; arrowheads, PROX1^+^ Ki67^−^ cells; dashed lines outline lymphatic vessels. Scale bar: 50 µm. One dot plot shows the percentage of PROX1^+^ cells that are Ki67^+^ (n = 4 mice), and the other shows Ki67^+^ PROX1^+^ cells in LD tissues. Data are shown as mean ± SD; differences were assessed using Student’s t-test.

To determine whether AQP1 upregulation is a common feature of inflammatory lymphangiogenesis beyond the *Pik3ca*^H1047R^ model, we examined its expression in a mouse model of secondary lymphedema (65). In this model, hindlimb lymphatic damage is induced through a combination of local irradiation of the inguinal region and surgical ablation of the inguinal lymph node, along with ligation of adjacent lymphatic vessels, leading to impaired lymphatic drainage, vessel dilation, and remodelling, tissue fibrosis and inflammation (65,66). Lymphatic injury triggers an inflammatory response, involving macrophages, T-helper and CD8^+^ T cells, neutrophils, and pro-inflammatory cytokines and lipids, such as TNFα, and LTB₄, which drive both acute and chronic inflammation (67). As expected, we observed a significant increase in CSF1R^+^ macrophages (68) and CD4^+^ T cells (69) in lymphedematous skin compared to controls (Figure S4D). Interestingly, similar to lymphedematous skin, CSF1R^+^ macrophages were abundant in the intestinal villi, where they were in direct contact with AQP1^+^ lacteals (Figure S4E). In contrast, macrophages were sparse and remained detached from AQP1^−^ submucosal lymphatic vessels.

Immunofluorescence analysis revealed markedly dilated VEGFR3^+^ lymphatic vessels in lymphedematous skin while only thin and collapsed lymphatic vessels were present in control samples (Figure 4D). Strikingly, AQP1 expression was highly and significantly increased in dilated lymphatic vessels in lymphedema, whereas AQP1 was undetectable in VEGFR3^+^ lymphatics of normal skin. AQP1-high and AQP1-low areas were detected within the same lymphatic vessel, suggesting the existence of local cues inducing its expression (Figure 4D). Additionally, uniform AQP1 upregulation was observed in EMCN^+^ blood vessels within lymphedema tissues (Figure S4F). To address the potential function of lymphatic AQP1 in lymphedema, we analyzed the correlation between its expression and LEC proliferation. Immunostaining for AQP1, PROX1 and proliferation marker Ki67 revealed an absence of proliferation in control skin samples and pronounced overall increase in Ki67^+^ LECs and stromal cells in lymphedema samples (Figure 4E). Most importantly, the proportion of proliferating Ki67^+^ cells was significantly higher in AQP1^+^ LECs as compared to AQP1^−^ population, suggesting that AQP1 may be important for lymphatic vessel remodelling under inflammatory conditions. Collectively, our findings in lymphatic malformations, secondary lymphedema and lymph node demonstrate that AQP1 expression is specifically induced during proinflammatory postnatal lymphatic remodelling but not during embryonic lymphangiogenesis.

## Discussion

In this study we identified three subpopulations of murine intestinal LECs, including lacteal, S/S (submucosal and serosal), and IFN subtypes, defined by characterizing markers and highlighting AQP1 expression as a specialized lacteal adaptation. Comparison of intestinal LECs with LECs from other organs shed new light on lacteal LECs, demonstrating their close similarity to *Ptx3*⁺ inflammatory LECs, while S/S LECs bear transcriptional traits characteristic of pre-collector lymphatics, allowing a clearer understanding of their distinct roles.

Lacteal LECs experience fluctuating osmotic conditions due to nutrient absorption, with tip osmolarities reaching approximately 600 mOsm (17). During fat absorption, the accumulation of chylomicrons and increased solute concentration in the lamina propria generate osmotic gradients, acutely altering lacteal junction configuration to allow chylomicron and nutrient entry (70). These conditions require rapid water movement; therefore, we propose that AQP1 is essential for swift adaptation of LECs to osmotic changes (71). Accordingly, *Aqp1* deletion in LECs caused lacteal shortening and decreased fat absorption, matching prior findings for shorter lacteals (16,17,72). This suggests that AQP1 plays a critical role in maintaining lacteal integrity, potentially by facilitating LEC migration through volume regulation necessary for protrusion formation (46,73,74). Supporting this notion, overexpression of *AQP1* enhanced LEC migration in a spheroid sprouting assay under hyperosmotic conditions in the presence of VEGF-C. The reduced lacteal length and diminished dietary lipid absorption observed in mice with LEC-specific *Aqp1* deletion likely explain their attenuated weight gain on HFD. Similar metabolic phenotypes were previously reported in mice with germline deletion of *Aqp1*, which showed reduced weight gain on HFD and decreased serum TG, with body weight normalizing on low-fat diet (29).

We observed a dynamic pattern of AQP1 expression in lacteals, with its intensity increasing after birth and peaking at the lacteal tip around P5. This timing coincides with the onset of milk intake (50) and a sharp increase in intestinal bacterial load (53), both likely contributing to osmolarity fluctuations. Concurrently, immune cells gather in the lamina propria, providing a source of VEGF-C and further influencing osmotic conditions. For instance, F4/80⁺ macrophages, initially concentrated at the villus base in 2-day-old neonates, expand dramatically over the next 12 days and ultimately reside predominantly in close proximity to capillaries, with occasional presence in the submucosa (75). Increased macrophage density parallels lacteal growth, underscoring how immune-derived VEGF-C fosters functional lacteals. Alongside smooth muscle cells and fibroblasts, these macrophages constitute major VEGF-C sources in the gut (17,72,76). Gut microbiota also shape the distribution and morphology of intestinal CX3CR1⁺ macrophages, with antibiotic treatment reducing their presence in the lamina propria but not in the submucosa (77). Consistent with the essential role of VEGF-C/VEGFR3 signalling in lacteal development and maintenance (16,17,50) both antibiotic treatment and VEGFR3 blockade diminished AQP1 expression. Reduced AQP1 levels under antibiotic treatment likely reflect lower VEGF-C in the lamina propria, aligning with evidence that gut microbiota modulates VEGF-C levels via macrophage-dependent mechanisms (76). Nevertheless, absence of AQP1 expression during VEGFR3-VEGF-C-dependent embryonic lymphangiogenesis suggests that additional immune-derived signals or osmotic factors are necessary for robust AQP1 induction.

To further contextualize our findings, lacteal LECs showed highest similarity with the *Ptx3*⁺ LEC subset, with elevated expression of *Ptx3*, *Itih5*, and *Mrc1*, genes associated with immune cell interactions. Specifically, MRC1 binds CD44^+^ lymphocytes (78), PTX3 increases responses to inflammatory signals (79), and ITIH5 interacts with PTX3 (80). This immune-interactive profile aligns with the known biology of lacteals, which are surrounded by macrophages in the lamina propria (Figure S4E, 75) and are continuously exposed to damage- and pathogen-associated molecular patterns derived from food and microbiota (15). Similarly, PTX3⁺ dermal LECs expand under inflammatory lymphangiogenesis and interact with macrophages (5). In lymph nodes, *Ptx3*⁺ LECs populate the medullary sinus, a region rich in F4/80⁺ macrophages (81) and dense with lymphocyte-macrophage clusters following immunization (82). These similarities suggest that local immune cues and tissue-specific signals may contribute to the emergence of specialized LEC states, enabling them to adapt to the unique settings of different tissues.

The similarity to immune-interacting LECs led us to explore the role of AQP1 in inflammatory dermal conditions. We previously reported that AQP1 is strongly induced in mesenteric LECs in a mouse model of human hereditary lymphedema-distichiasis (7). This study shows that AQP1 is markedly induced in LECs in secondary lymphedema and lymphatic malformations driven by hyperactive PI3K signalling (*Pik3ca^H1047R^*). During inflammatory lymphangiogenesis, macrophages secrete lymphangiogenic factors such as VEGF-C (83,84), potentially promoting both lymphatic vessel proliferation and AQP1 upregulation. Elevated AQP1 levels correlate with increased proliferative activity in these inflamed vessels, suggesting that AQP1 may drive, or at least mark, remodelling processes in lymphedema and lymphatic malformations. In contrast, embryonic dermal lymphangiogenic vessels exhibit robust sprouting and proliferation without AQP1 induction, indicating that while embryonic LECs rely on VEGF-C/VEGFR3 signalling, they lack the inflammatory, metabolic or osmotic cues needed for AQP1 upregulation. Moreover, embryonic macrophages are not the primary drivers of lymphangiogenesis in the developing skin (85), further emphasizing the contextual nature of AQP1 expression. Consistent with this, AQP1 protein was not detected in normal dermal LECs, despite the presence of *Aqp1* transcripts, possibly due to additional posttranscriptional mechanisms that regulate AQP1 protein stability. One explanation is that AQP1 function is particularly relevant under the biophysical stresses associated with adult inflammatory states, rather than during tightly regulated embryonic development. Indeed, osmolarity rises in lymphedema (86), and concurrent AQP1 induction may either spur or reflect excessive vessel growth. Because AQP1 is generally absent where osmotic demand is low and baseline membrane permeability of water suffices, contextual increases in osmotic stress, such as those seen in the small intestine or lymphedema, may also contribute to AQP1 expression to accommodate rapid water flux and enhanced cell migration.

Collectively, our findings establish AQP1 as a hallmark of actively remodelling postnatal LECs within inflammatory microenvironments. We identify a beneficial role for AQP1 in LEC migration under high osmolarity conditions, offering insight into the pathophysiological mechanisms of lymphedema and potential therapeutic interventions. Inflammation promotes the accumulation of extracellular proteins and Na⁺, thereby elevating tissue osmolarity (87). Although this short-term increase in osmolarity supports immune cell activation and aids in combating infections, we propose that in lymphedema, chronic inflammation coupled with defective lymphatic drainage raises osmolarity to a level that impedes regenerative lymphangiogenesis, even in the presence of abundant VEGF-C. Thus, strategies aimed at reducing pathological osmolarity may complement lymphangiogenic growth factor therapies.

## Materials and methods

### Animal studies

All animal care and experimental procedures were performed in accordance with relevant national and institutional guidelines, following approval by the Animal Ethics Committee of Vaud (Switzerland), the Uppsala Animal Experiment Ethics Board (Sweden), or the local Animal Ethical Committee at the University of Liège (Belgium). Mice were on C57BL/6J background and were maintained under specific pathogen-free conditions with a 12-hour light/dark cycle and *ad libitum* access to food and water. Unless otherwise specified, experiments were conducted using age- and sex-matched cohorts of adult mice aged 8–12 weeks.

#### Generation of inducible Aqp1 knockout mice

To generate tamoxifen-inducible, LEC-specific *Aqp1* knockout mice (*Aqp1^iΔLEC^*), *Aqp1^fl/fl^* mice (48) were crossed with *Prox1-CreERT2* (49) mice. To induce *Aqp1* deletion, tamoxifen (T5648, Sigma-Aldrich) was dissolved by sequentially adding 100% ethanol, Kolliphor EL (Sigma-Aldrich, Cat# C5135), and phosphate buffer saline (PBS) in a 1:1:8 ratio, with thorough mixing after each addition, to prepare a stock solution at a concentration of 10 mg/ml. Control *Aqp1^fl/fl^* and *Aqp1^iΔLEC^*mice received intraperitoneal injections of tamoxifen at a dose of 50 μg/g body weight. Injections were administered three times every other day during the first week, followed by one injection per week until sacrifice.

#### R26-Pik3ca^H1047R^; Vegfr3-CreERT2 mice

were generate by crossing of *R26-Pik3ca^H1047R^* (88) with *Vegfr3-CreERT2* mice (89). Dermal lymphatic malformations were induced as previously described (63). Briefly, 50µg of 4-hydroxy-tamoxifen dissolved in acetone was applied topically to the ear of 3-week-old mice. Mice were sacrificed 3 weeks later, and the dorsal ear skin was dissected from underlying cartilage and fixed in 4% (paraformaldehyde) PFA in PBS for 2 hours at room temperature. Tissue was subsequently washed twice with PBS before proceeding to immunostaining.

#### Secondary lymphedema model

The hindlimb lymphedema model combined local irradiation and surgical intervention as previously described (65). Lymphedema was induced in the left limb, with the right limb serving as an internal control. For irradiation, mice were anesthetized with 2% isoflurane and positioned ventrally inside a precision X-ray irradiator. To accurately target the inguinal region, X-ray radiography (40 kV, 0.5 mA) was performed. A 20 mm-square collimator was placed over the targeted area to deliver a single 30 Gy dose (225.0 kV, 13.00 mA) in an anteroposterior direction. The left limb was irradiated for 317 seconds on the ventral side and an additional 317 seconds on the dorsal side. One week later, surgery was performed under 2% isoflurane anesthesia on a 37°C heating pad within a horizontal airflow hood. The left hindlimb was shaved, disinfected with dermal isobetadine, and injected with 5 µL of 2% Evans Blue dye between the footpads to visualize lymphatic structures. Lymphatic injury was induced through a three-steps surgery: i) a circumferential skin incision at the inguinal level, ii) excision of the inguinal and popliteal lymph nodes, and iii) ligation of collecting lymphatic vessels parallel to the ischial vein with three separate 7/0 non-absorbable polypropylene sutures under a 10x binocular magnifier. Finally, the skin was sutured with 5/0 non-absorbable silk stitches.

#### Antibiotic treatment

Wild-type *Aqp1^fl/fl^* and *Aqp1^iΔLEC^* mice were co-housed from weaning to minimize microbiota differences. Three weeks after tamoxifen administration, adult mice were treated with a broad-spectrum antibiotic cocktail in their drinking water for four weeks. The treatment consisted of enrofloxacin (2.5 mg/ml; Baytril 10%, Bayer) for the first two weeks, followed by amoxicillin (0.8 mg/ml) and clavulanic acid (0.114 mg/ml; Co-Amoxi-Mepha, Mepha) for the subsequent two weeks (7). Control mice received regular drinking water.

#### High-fat diet (HFD) feeding

After inducing *Aqp1* deletion with tamoxifen over a two-week period - three doses administered every other day during the first week and one dose during the second week - the mice were fed either a HFD (D12492i, Research Diets: 60 kcal% fat, 20 kcal% protein, 20 kcal% carbohydrate) or a matched normal chow diet (NCD; D12450Bi, Research Diets: 10 kcal% fat, 20 kcal% protein, 70 kcal% carbohydrate) for three weeks. Mice were randomly assigned to either the HFD or NCD group, and diets were provided ad libitum. Body weights were recorded weekly.

#### VEGFR3 signalling blockade

Neonatal mice were administered subcutaneous injections of the VEGFR3-blocking antibody mF4-31C1 (54) or isotype-matched IgG2a antibody at P4, P6, and P8 at dose of 40 μg/g body weight using 30G needle. Tissues were harvested on P10 for analysis.

#### Lipid absorption assay

Mice were fasted overnight and then orally gavaged with olive oil (O1514, Sigma-Aldrich) at a dose of 10 μl/g body weight using a 20G, 60 mm gavage needle. Blood samples were collected from the tail vein at baseline (0 hours) and at 2- and 4-hours post-gavage into lithium heparin-coated tubes (Microvette CB 300 Hep-Lithium, Sarstedt). Samples were kept on ice for 30 minutes and centrifuged at 14,000 × g for 15 minutes at 4 °C to obtain plasma, which was stored at – 80 °C until analysis. Plasma triglyceride levels were measured on diluted samples (1:1 plasma to diluent ratio) using the Dimension® Xpand Plus system (Siemens Healthcare Diagnostics AG, Düdingen, Switzerland) according to the manufacturer’s protocol for triglyceride measurement (Siemens Healthcare, DF69A).

#### Lymph node analyses

Embryonic (E17.5-E18.5) and young adult male (9 weeks) inguinal lymph nodes were used for control stainings. For immunization, 16-17-weeks old females were injected with oligodeoxynucleotides CpG (Microsynth) and keyhole limpet hemocyanin (NP-KLH) (10 µg each in 10 µl) into the footpads. On day 7 mice were sacrificed, and inguinal lymph nodes were harvested and processed for cryosections.

### Generation of *AQP1*-overexpressing LECs

Human intestinal lymphatic endothelial cells (hLECs) were isolated from discarded surgical specimens with informed consent and institutional approval and cultured in EBM™-2 Endothelial Cell Growth Basal Medium-2 (EBM-2) medium (Lonza, CC-3156) supplemented with growth factors (EGM-2 Endothelial SingleQuots Kit, Lonza, CC-4176) and 1% fetal bovine serum (FBS) at 37°C in a humidified atmosphere with 5% CO₂.

To establish a stable LEC line overexpressing human *AQP1* (*AQP1*^OE^ LECs), cells were transduced with lentiviral particles containing human *AQP1* cDNA under the EF1α promoter (pLV[Exp]-Puro-EF1A>FLAG/hAQP1[NM_198098.4], VectorBuilder). Control cells were transduced with lentiviral particles encoding enhanced blue fluorescent protein (*EBFP*). Briefly, LECs were seeded in 6-well plates (0.2 × 10^6^ cells/well) when they reached 70% confluency, they were first starved for FBS and growth factors for 2 hours, then incubated overnight with lentivirus at a multiplicity of infection 15 (MOI). The virus-containing medium was replaced with complete medium the following day. Stable cell lines were selected using puromycin (350 ng/ml; Sigma-Aldrich, P7255) for 3-5 days. Transduction efficiency of *AQP1* was confirmed by fluorescence microscopy and western blot.

### Spheroid sprouting assay

The spheroid sprouting protocol described below was adapted from (90). LECs were suspended at 800 cells per spheroid in a mixture of 72% EBM-2 complete medium, 8% FBS, and 20% methylcellulose stock solution (2% methylcellulose in Medium 199 with GlutaMAX™). Aliquots of 100 μl were dispensed into each well of non–tissue culture-treated 96-well round-bottom plates and incubated overnight at 37 °C with 5% CO₂ to form spheroids.

After 20 hours, 5-7 spheroids were collected and mixed in a fibrin gel composed of 2.5 mg/ml fibrinogen (Sigma, F8630) in EBM-2 complete medium, supplemented with aprotinin (final concentration 0.1 U/ml, Sigma, A1153) and either bovine serum albumin (BSA) or VEGF-C at 100 ng/ml. Gelation was initiated by adding 30 µl thrombin (final concentration 0.625 U/ml; Sigma, T9549) to the spheroid-fibrinogen mixture (570 ul) in pre-warmed 24-well plates. The mixture was gently mixed and incubated for 10-15 minutes at room temperature and 37 °C for 45 minutes. After gel formation, 600 μl of EBM-2 medium with 30,000 fibroblasts was added per well, and plates were incubated for 48 hours at 37°C with 5% CO₂. Spheroids were then fixed and stained with phalloidin and Hoechst for imaging. To minimize variability, spheroids for all conditions within each replicate were embedded in the same 24-well plate using the same batch of fibrin gel. After 48 hours, spheroids were fixed with 4% PFA for 15 minutes, permeabilized with 0.3% Triton X-100 in PBS for 10 minutes, and stained with Alexa Fluor 488-phalloidin (1:400; Invitrogen) and Hoechst dye (1 μg/ml; Sigma-Aldrich) to visualize F-actin and nuclei.

Normosmotic, hyperosmotic, and hypotonic conditions were established by adjusting the osmolality of the added culture medium (600 μl). Normosmotic medium was complete EBM-2 (approximately 300 mOsm/kg). Hyperosmotic medium was prepared by adding D-sorbitol (200 mM; Sigma-Aldrich, S1876), increasing the osmolality by 200 mOsm/kg. Hypotonic medium was made by 30% dilution complete EBM-2 with sterile distilled water.

Spheroids were imaged with a Leica Ti2 confocal laser scanning microscope equipped with a 20× objective. Images were analyzed in ImageJ by measuring the distance from the spheroid core edge to the sprout tip. For each condition, 3-6 spheroids were assessed across four replicates. The mean sprout length per spheroid was calculated and normalized to the control condition (EBFP^OE^, normosmotic, BSA-treated).

### Isolation of intestinal mouse LECs

#### Tissue processing

Small intestinal segments (excluding the duodenum) were collected using curved scissors and washed by gently vortexing in 50 ml Falcon tubes containing cold PBS, followed by a second wash in cold PBS. To remove epithelial cells, the tissues were incubated at 37 °C and 150 rpm in Hank’s Balanced Salt Solution (HBSS) containing 5 mM EDTA, 1 mM DTT, and 20 μM HEPES for 15–20 minutes, briefly vortexed, rinsed in HBSS until the supernatant was clear, and then enzymatically digested in HBSS containing Liberase TL at a final concentration of 0.9 U/ml and DNase I at 0.1 mg/ml for 60 minutes at 37 °C, 180 rpm. Every 20 minutes, tissues were pipetted up and down to promote dissociation; the resulting supernatant was passed through a 70 μm strainer into 10 ml cold HBSS with 10% FBS. The remaining tissue fragments were returned to fresh digestion medium. After three rounds of digestion, cell suspensions were centrifuged at 1200 rpm for 5 minutes at 4 °C.

For scRNAseq experiments, we obtained intestinal tissues from three female and two male C57Bl/6 mice (9-10 weeks old), whereas for qPCR analyses we utilized samples from three *Aqp1^fl/fl^* and three *Aqp1^iΔLEC^* mice.

#### Cell staining and FACS sorting

Cells were resuspended in PBS containing 5% FBS and incubated with anti-CD16/32 (2.4G2) hybridoma supernatant to block non-specific IgG binding. Cells were stained with conjugated antibodies: for scRNA-seq, CD45-FITC, CD31-BV421, and PDPN-PE; for qPCR, EpCAM-BV421, CD45-PE-Cy7, CD31-PE, and PDPN-APC. Before sorting, 0.1 μg/ml DAPI and RedDot1 were added for 5 minutes at room temperature to exclude dead cells and debris. LECs were gated as DAPI⁻ redDot⁺ CD45⁻ CD31⁺ PDPN⁺ cells. For qPCR samples, an additional exclusion of EpCAM⁺ cells (EpCAM⁻) was applied. Cells were sorted into HBSS containing 30% FBS at 4 °C on a Beckman Coulter MoFlo Astrios EQ cell sorter equipped with a 70 µm nozzle, operating at 60 psi. The sort precision was set to “enrich” to maximize yield. On average, 23,000 LECs were sorted per sample, and sorting experiments were performed in three independent replicates.

### Validation of *Aqp1* deletion via RT-qPCR

Total RNA was isolated using the RNeasy Micro Kit (Qiagen). The concentration and quality of the isolated RNA were assessed using an Agilent 2100 Bioanalyzer. Reverse transcription was performed using the Transcriptor First Strand cDNA Synthesis Kit (Roche Diagnostics, Cat# 04379012001). Prior to cDNA synthesis mRNA was amplified using the Ovation Pico WTA System V2 (NuGEN).

Quantitative real-time PCR (RT-qPCR) analyses were conducted using the StepOnePlus Real-Time PCR System (Applied Biosystems) with SYBR Green PCR Master Mix (Thermo Fisher Scientific). Primers flanking exons 2 and 3 of the *Aqp1* gene (48) were employed to confirm the deletion of this region in tamoxifen-induced *Aqp1^iΔLEC^* samples. The housekeeping gene 18S was used as an internal control, and results were presented as relative expression levels normalized to control samples.

### Western Blot analysis

Cell lysates were prepared using RIPA buffer with protease inhibitors (Roche, 11697498001). Protein concentration was measured using the BCA assay (Pierce, 23227). Proteins were separated by SDS-PAGE, transferred to PVDF membranes, blocked with 5% bovine serum albumin (AppliChem, A1391) and 0.3% Tween 20 (AppliChem, A1389) in PBS, and probed with antibodies against AQP1 and Histone H3. HRP-conjugated secondary antibodies were used, and signals were detected using an ECL substrate (GE Healthcare).

### Immunohistochemistry and imaging

#### Tissue preparation and staining

Following euthanasia, the mice were intracardially perfused first with PBS and then with 4% PFA (158127, Sigma-Aldrich) to fix the tissues. Preparation of whole-mounted tissues, cryosections, and paraffin sections were performed as previously described in (91). In brief, the small intestine was dissected between the stomach and cecum, flushed with cold PBS, opened longitudinally, and pinned flat. Tissues were fixed in 4% PFA (paraffin sections) or 0.5% PFA with picric acid and sodium phosphate (whole-mounts and cryosections) overnight at 4°C. Cryosections were cut at 10 μm (gut) or 8 μm (LN), and paraffin sections at 4 μm. Sections were permeabilized with 0.3% Triton X-100, blocked with 0.5% BSA, and incubated with primary antibodies overnight at 4°C. After PBS washes, sections were stained with Alexa Fluor-conjugated secondary antibodies for 1 hour (sections) or overnight (whole-mounts). DAPI was added for nuclear staining. Samples were mounted in Fluoromount-G (sections) or Histodenz (whole-mounts).

#### Multiplex immunohistochemistry

Multiplex immunohistochemistry was performed on paraffin-embedded tissue sections. After the initial round of primary antibody staining and imaging, coverslips were removed by incubating the sections in PBS 1x with gentle rocking and sections were incubated with elution buffer composed of 0.5 M glycine, 3 M guanidinium chloride, 3 M urea and 40 mM tris(2-carboxyethyl) phosphine in distilled water for 4 min at RT. Following elution and washes, sections were incubated with a second set of primary antibodies for a second round of staining.

#### Immunostaining of cells

LECs seeded on glass coverslips in 24-well plates, were fixed with 4% PFA in PBS containing 0.1% CaCl₂ and 0.05% MgCl₂, permeabilized with 0.1% Triton X-100 (AppliChem, A1388) in PBS and blocked with a solution of 0.5% BSA (AppliChem, A1391) and 0.1% Triton X-100 in PBS for 1 hour at room temperature. Coverslips were incubated with primary antibodies incubated overnight at 4 °C, washed with PBS, incubated with Alexa Fluor-conjugated secondary antibodies (Thermo Fisher Scientific) supplemented with DAPI (0.2 μg/ml) for 1 hour on a shaking platform at room temperature. Following incubation, coverslips were washed and mounted using Fluoromount-G mounting medium.

#### Imaging and analysis

Confocal imaging was performed using the following systems and software: Zeiss LSM 880 with Airyscan (Zen Black software), Nikon Ti2 Yokogawa CSU-W1 Spinning Disk (Nikon NIS-Elements AR 5.0 software), Zeiss Imager Z1 (Zeiss Axiovision SE64 rel 4.9.1 software), and Hamamatsu NanoZoomer S60 (NDP.view2 software). We used ImageJ Fiji for image analysis. To quantify AQP1 and VEGFR3 co-expression, a threshold was first set on the VEGFR3 channel to delineate the total lymphatic vessel area, followed by a separate threshold for the AQP1 channel. The area of double-positive (AQP1^+^VEGFR3^+^) regions was then measured and divided by the total VEGFR3^+^ area. The same approach was used to determine the percentage of AQP1^+^EMCN^+^ area. For LEC proliferation, thresholds were set for both PROX1 and KI67, and the number of double-positive PROX1^+^KI67^+^ cells was normalized to the total number of PROX1^+^ cells. A threshold based on the Otsu algorithm was applied to the AQP1 channel to classify proliferating LECs (PROX1^+^KI67^+^) as AQP1^+^ or AQP1^−^, based on whether the measured mean AQP1 intensity in each cell exceeded this threshold. Macrophages were quantified by applying a threshold to the CSF1R channel and counting the number of CSF1R^+^ cells, which was subsequently normalized to the total tissue area.

### Single-cell transcriptomic analysis

#### scRNA-seq library preparation, sequencing and pre-processing

Sorted LECs (∼10,000 cells per sample) were loaded into the Chromium Controller (10x Genomics) using the Single Cell 3’ Library & Gel Bead Kit v3. Libraries were prepared following the manufacturer’s protocol. Paired-end sequencing was performed on Illumina HiSeq 2500 (150 cycles) and Illumina NovaSeq 6000 (100 cycles) devices at the Lausanne Genomics Technology Facility to achieve a depth of at least 50,000 reads per cell. Raw sequencing reads were aligned to the mm10 mouse reference genome (v. mm10-3.1-0 of the transcriptome reference provided by 10x Genomics), number of reads were summarized per gene, and cells were called using Cell Ranger software (v3.1.0, 10x Genomics).

#### Data acquisition and integration

Our scRNA-seq data (samples 1–5) was integrated with three publicly available datasets (samples 6-8). Raw count matrices of all datasets were imported into R v4.1.2 for further analysis. Each dataset was ln-normalized with a scale factor of 10,000, followed by variable gene detection using the Seurat package v4.3.0 (92). LECs were identified based on the expression of canonical markers (*Flt1*^−^, *Pecam1*^+^, *Prox1*^+^, and *Flt4*^+^). To ensure data quality, cells with mitochondrial gene expression exceeding 5% and dissociation-related genes (93) exceeding 6% were filtered out. Integration of retained cells was performed using the anchor correspondence method implemented in the Seurat package v4.3.0, using 30 dimensions and 60 nearest neighbours as input parameters (92,94).

#### Unsupervised clustering and visualization

Dimensionality reduction was performed using principal component analysis (PCA) on the top 2000 variable genes, followed by Uniform Manifold Approximation and Projection (UMAP) for visualization, using 25 principal components as input. Clustering was achieved using a shared nearest neighbour (SNN), Louvain modularity optimization-based clustering algorithm, with 25 principal components, 20 nearest neighbours and a resolution of 0.3 as input. Cell types were annotated based on the expression of canonical markers, identifying three distinct LEC populations.

#### Differential gene expression, pathway and similarity analysis

Differential gene expression within each cluster compared to all other cells was assessed using the FindAllMarkers function of the Seurat package, which implements a Wilcoxon rank-sum test with Bonferroni correction for multiple comparisons. Genes with adjusted p-values < 0.05 were considered significantly differentially expressed. Visualization tools within Seurat, such as dot plots and violin plots, were used to display gene expression patterns. Over-representation analysis of Gene Ontology (GO) Biological Process terms was performed by separating the significantly up-and down-regulated genes in each cluster with the enrichGO function of the clusterProfiler package v4.2.2 (95,96,97). We manually parsed the list of significant GO terms to remove terms that had redundant gene content, or shared ancestor terms, and calculated a module score of the genes of each selected GO term per cell, using the AddModuleScore (98) function of the Seurat package. The average module score per cluster per GO term was calculated and displayed as a heatmap using the ComplexHeatmap package v2.10.0 (99) for R.

We investigated whether intestinal LECs expressed gene signatures of LECs from other organs such as the mesentery, the skin (deriving either from control or *Pik3ca*^H1047R^ mutant mice) and the lymph node. To this end, we obtained transcriptomics data from the respective organs. For each mesenteric (7) or dermal LEC subtype (5), the FindAllMarkers function was used to define a signature consisting of 50 marker genes up-regulated in each LEC subtype. For lymph node LEC gene signatures, we used the marker genes described in Figure 5B from Xiang et al. (6). The AddModuleScore function was used to evaluate the expression magnitude of the mesenteric, dermal or lymph node LEC subtypes per cell, and the median module score per gene signature per cluster displayed using the ComplexHeatmap package. Finally, we assessed transcription factor activity per cell using the SCENIC implementation for R v.1.3.1, using default parameters (36).

### Statistical analysis

Data were analysed using Wolfram Mathematica (13.0), R (v4.1.2) and GraphPad Prism (8). All plotted values are presented as mean ± standard deviation (SD). For two-group comparisons, a one-sample, paired, Student’s or Welsh’s t-test was performed, as appropriate. For multiple group comparisons, two-way ANOVA or two-way repeated-measures ANOVA was used, followed by Tukey’s post hoc test or p-value corrections (Benjamini-Hochberg or Holm–Sidak). The specific test used for each experiment are indicated in the figure legends. A *p* value <0.05 was considered statistically significant.

---

**Supplementary to Figure 1.**
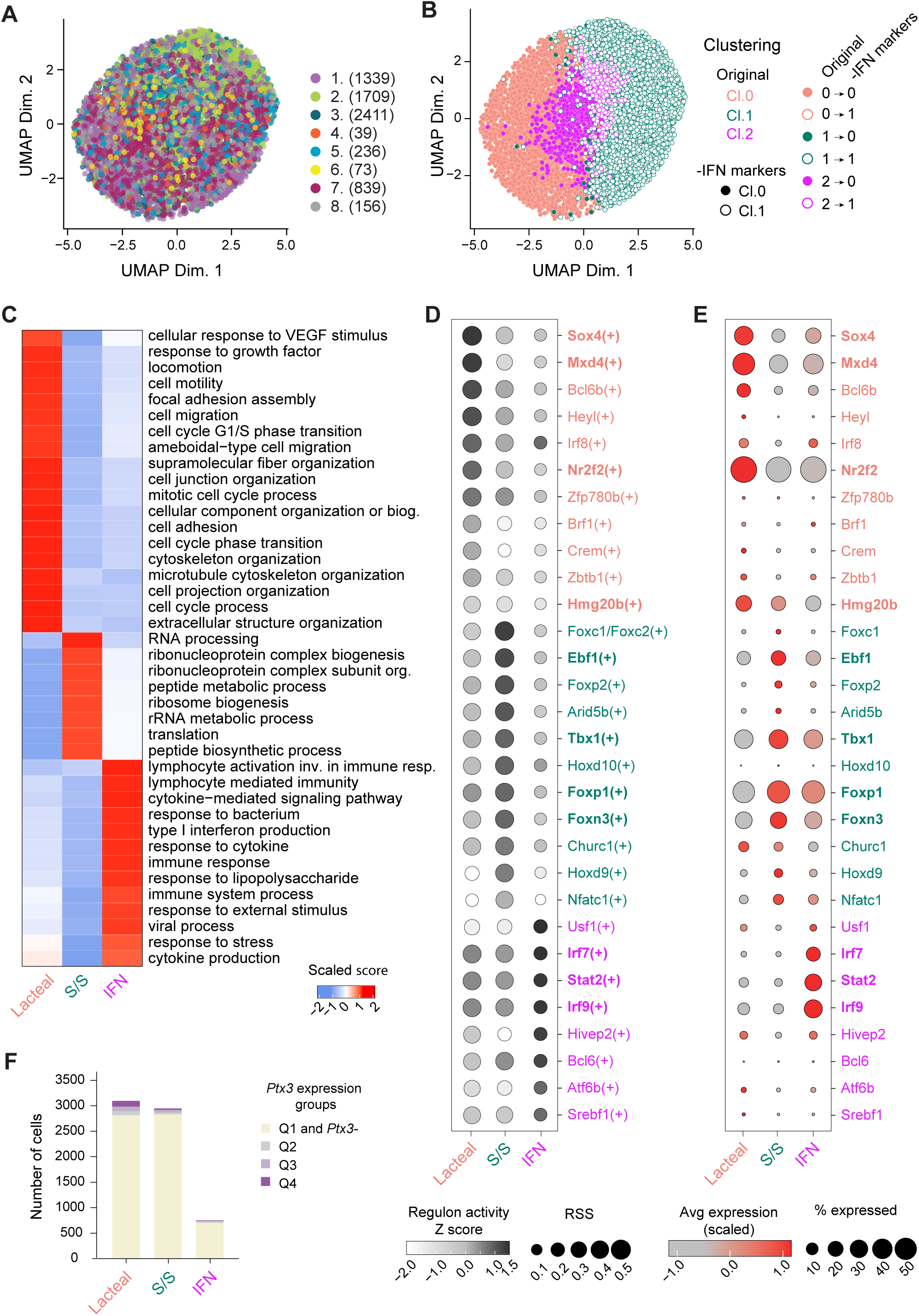
**A.** UMAP of all integrated cells colored by their originating sample (dataset numbers per Fig. 1A; cell numbers in brackets). **B.** UMAP of integrated cells re-clustered after removing cluster 2 markers. Colors indicate original clustering (Cl.0-1-2; Fig.1B), with full or empty circles distinguishing the re-clustering of Cl.0-1. **C.** Heatmap of top GO biological processes per cluster, colored by enrichment score. **D.** Dot plot of the top 10 transcription factors with highest variability for regulon activity among clusters. Dot size shows Regulon Specificity Score (RSS), indicating specificity to a cluster; dot color reflects the regulon activity z-score, indicating the level of activity. Transcription factors expressed in the respective clusters are in bold. **E.** Dot plot of the top 10 transcription factors with variable regulon activity. Dot size indicates the percentage of cells expressing each factor; color reflects scaled average expression. **F.** Bar chart of *Ptx3* expression levels in intestine LEC clusters. Colors indicate *Ptx3* expression quartiles: 0–25%, 25–50%, 50–75%, 75–100%.

**Supplementary to Figure 2.**
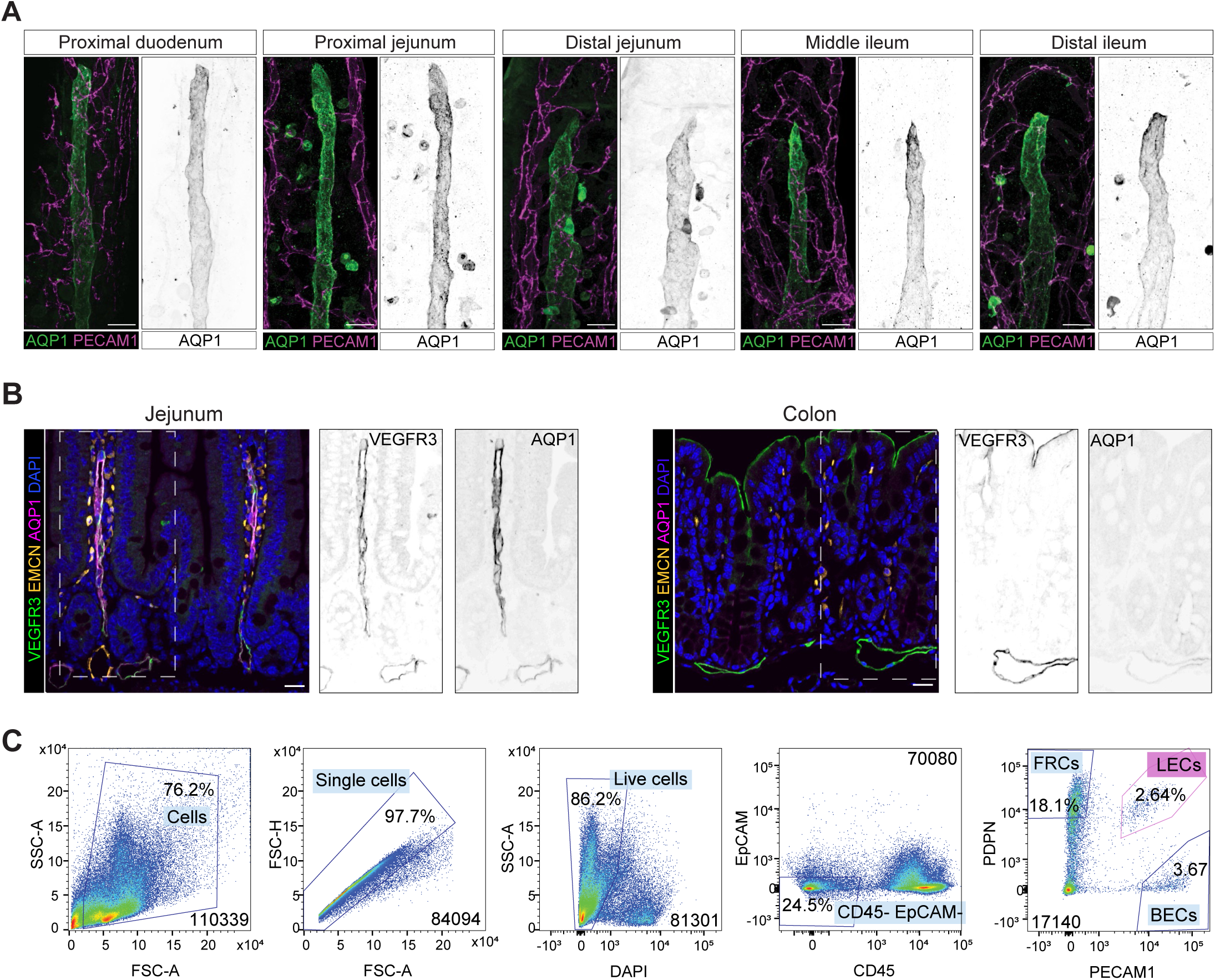
**A.** AQP1 is expressed in lacteals throughout the intestine. Duodenum, jejunum and ileum stained for AQP1 (green) and PECAM1 (purple); grayscale images show AQP1 alone. Scale bar: 25 μm. **B.** AQP1 is detected in intestinal but not colon LECs. Paraffin sections of jejunum and color stained for AQP1 (magenta), EMCN (yellow), VEGFR3 (green), DAPI (blue); grayscale images show individual markers for insets. Scale bar: 20 μm. **C.** Flow cytometry gating for LEC isolation from the small intestine. Gating steps include exclusion of debris and doublets, gating of DAPI^−^ live cells, exclusion of CD45^+^ and EpCAM^+^ cells, and finally sorting of PECAM1^+^PDPN^+^ LECs.

**Supplementary to Figure 3.**
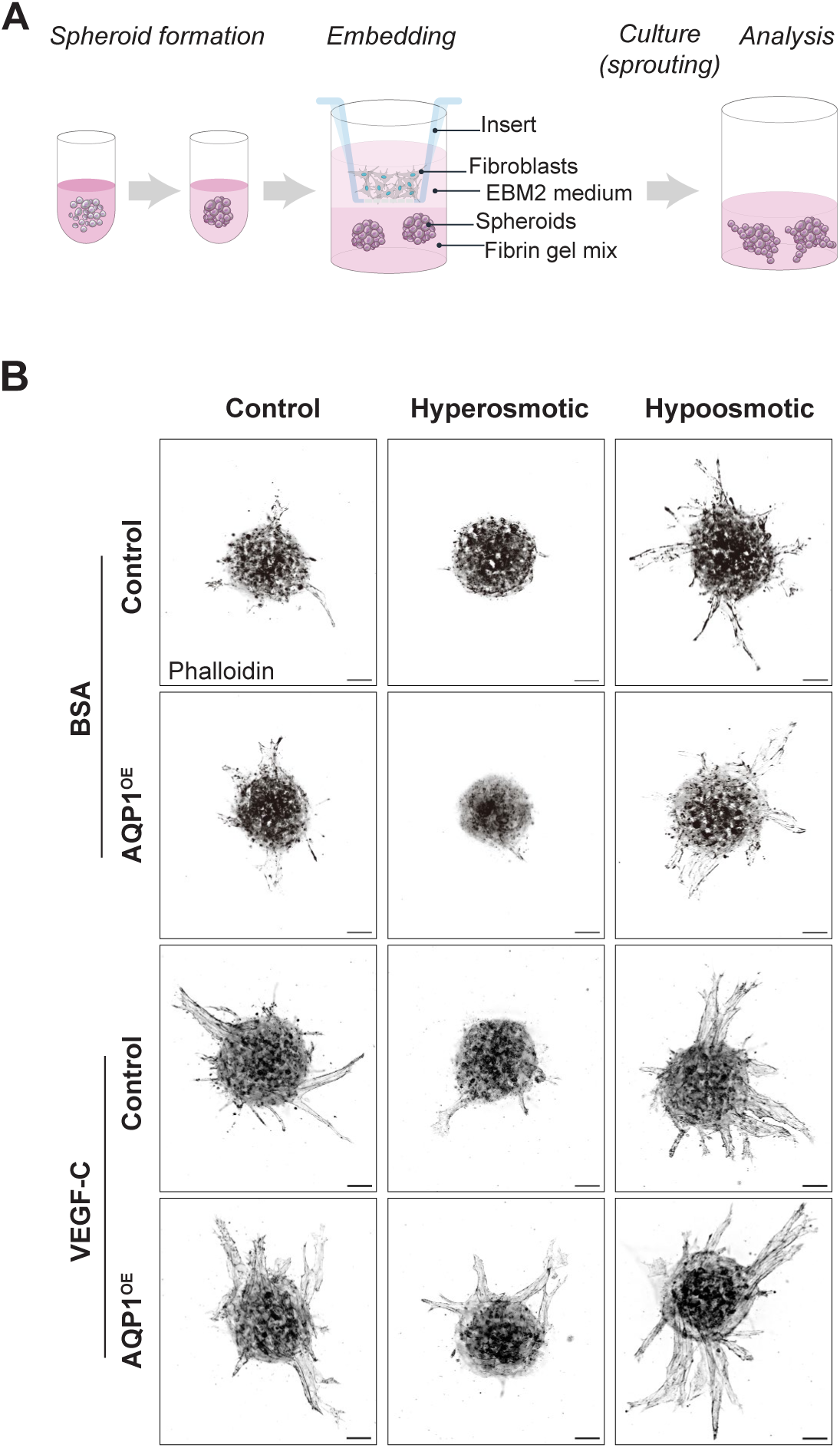
**A.** Scheme of the spheroid sprouting assay. **B.** Representative phalloidin-stained spheroids from control (*EBFP^OE^*) and *AQP1^OE^*LECs, cultured in normosmotic, hyperosmotic, or hypoosmotic medium, with or without VEGF-C. Scale bar: 50 µm

**Supplementary to Figure 4.**
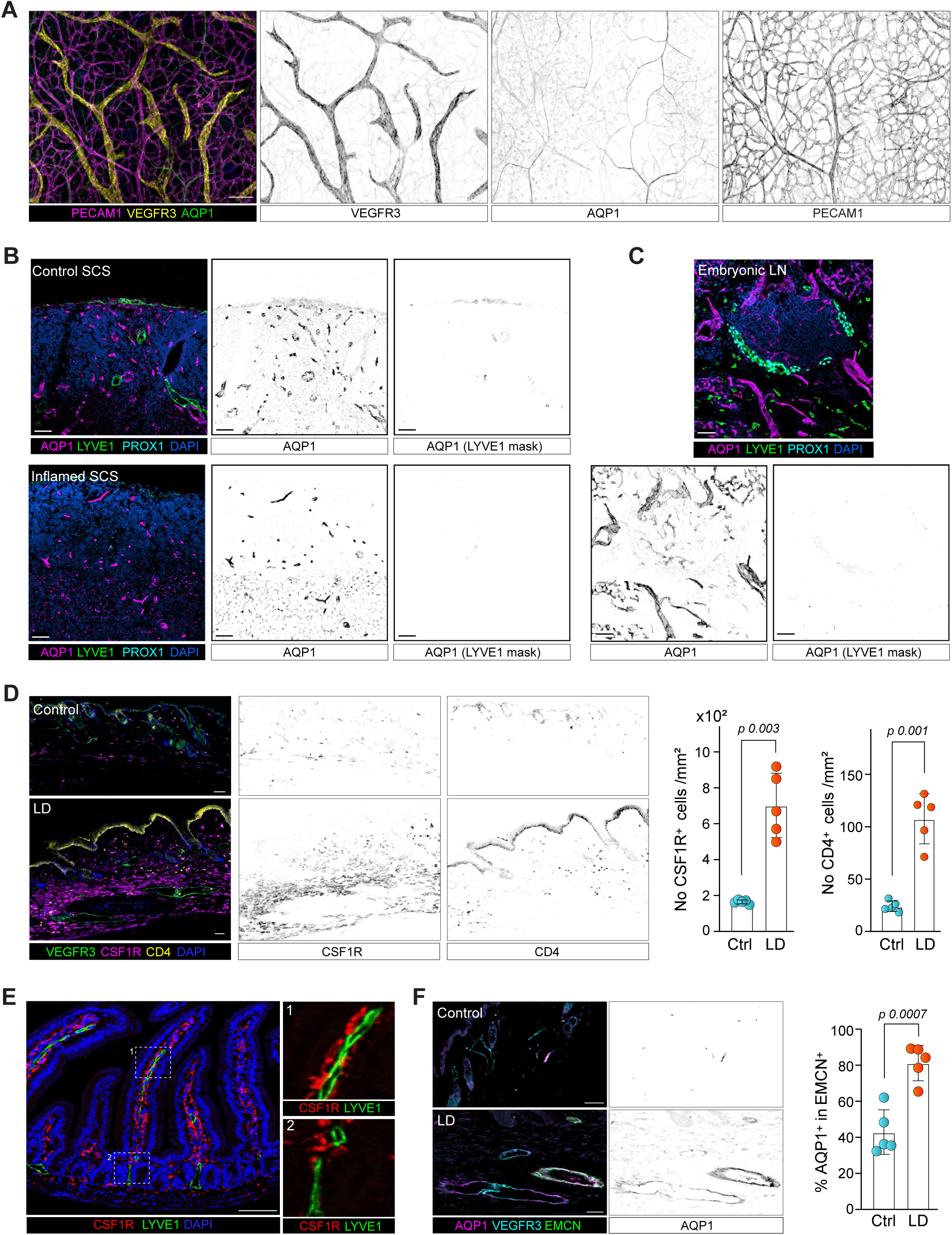
**A.** AQP1 is not expressed during embryonic lymphangiogenesis. Whole mount staining E16.5 skin stained for PECAM1 (magenta), VEGFR3 (yellow), AQP1 (green); grayscale images show individual markers. Scale bar: 100 µm. **B.** AQP1 is absent in subcapsular sinus LECs. Control and inflamed LN sections stained for AQP1 (magenta), LYVE1 (green), PROX1 (cyan), and DAPI (blue); grayscale images show AQP1 alone and AQP1 masked by LYVE1 staining. Scale bar: 50 µm. **C.** AQP1 is absent in embryonic LN LECs. E17.5 LN sections stained for AQP1 (magenta), LYVE1 (green), PROX1 (cyan) and DAPI (blue); grayscale images show AQP1 alone and AQP1 masked by LYVE1 staining. Scale bar: 50 µm. **D.** Macrophage infiltration increases in lymphedema (LD). Skin sections (control and LD) stained for VEGFR3 (green), CSF1R (magenta, macrophages), CD4 (yellow, T-cells), and DAPI (blue); grayscale images show CSF1R or CD4 alone. Scale bar: 50 µm. The dot plot quantifies the number of CSF1R^+^ cells per mm². **E.** AQP1^+^ lacteals are surrounded by macrophages. Paraffin sections of jejunum stained for LYVE1 (green), CSF1R (red), and DAPI (blue) (n=5 mice). Scale bar: 20 µm. **F.** AQP1 is upregulated in blood vessels under LD. Skin sections (control and LD) stained for AQP1 (magenta), VEGFR3 (cyan), and EMCN (green); grayscale images show AQP1 alone (n=5 mice). Scale bar: 50 µm. The dot plot shows the percentage of AQP1^+^ area within EMCN^+^ vessels. Data are mean ± SD. Differences were assessed using Student’s t-test.

## Acknowledgements

We thank Vedat Schwenger and Andreas Wagner for providing the *Aqp1^fl/fl^* mice, Kari Alitalo for generously sharing VEGF-C and Werner Held for providing anti-CD16/32 (2.4G2) hybridoma supernatant. We are grateful to Céline Beauverd for mouse genotyping and her assistance with immunohistochemistry. We gratefully acknowledge the Mouse Pathology, Flow Cytometry, Genomic Technologies Facility (GTF), Animal, and Cellular Imaging Facilities at the University of Lausanne for their assistance. This work was supported by the European Union’s Horizon 2020 research and innovation program Theralymph (grant agreement no. 847939) to TVP, IR, AN and TM; the Swiss National Science Foundation (310030_197878) to TVP; Human Frontier Science Program (LT000074/2019-L) to JK; the Muschamps Foundation and the Swiss National Science Foundation (CRSK-3_190435) to JBL; Human Frontier Science Program (LT000633/2020-L) to SA-M; European Union’s Horizon 2020 research and innovation programme under the Marie Skłodowska-Curie grant agreement No 814316 to HS and TM; and the Fonds de la Recherche Scientifique-FNRS (F.R.S.-FNRS, Belgium) to AN. The authors have no conflicts of interest to declare.

## Notes

### Competing Interest Statement

The authors have declared no competing interest.

